# Coherent waves of myelin plasticity during motor-skill learning

**DOI:** 10.1101/2021.04.13.439402

**Authors:** Michela Azzarito, Gabriel Ziegler, Eveline Huber, Patrick Grabher, Martina F. Callaghan, Alan Thompson, Karl Friston, Nikolaus Weiskopf, Tim Killeen, Patrick Freund

**Author notes:** Correspondence to: Dr Patrick Freund; Spinal Cord Injury Center Balgrist; University Hospital Zurich, University of Zurich, Forchstrasse 380, 8008 Zurich, Switzerland.

## Abstract

Motor skill learning relies on neural plasticity in the motor and limbic systems. However, the spatial and temporal dependencies of these changes—and their microstructural underpinnings—remain unclear. Eighteen healthy males received training in a computer-controlled motion game 4 times a week, for 4 weeks. Performance improvements were observed in all trained participants. Serial myelin-sensitive multiparametric mapping at 3T during this period of intensive motor skill acquisition revealed temporally and spatially distributed, performance-related myelin-sensitive microstructural changes in the grey and white matter across the corticospinal system and hippocampus. Interestingly, analysis of the trajectory of these transient changes revealed a time-shifted choreography across white and grey matter of the corticospinal system as well as with changes in the hippocampus. Crucially, in the cranial corticospinal tracts, myelin-sensitive changes during training in the posterior part of the limb of the internal capsule were of greater magnitude in lower-limb trainees compared to upper limb trainees. Motor skill learning is depended on coherent waves of plasticity within a corticospinal-hippocampal loop.

## Introduction

Motor skill learning of a complex sensorimotor task, such as juggling or dancing, requires physical and cognitive effort—and relies on functional changes within the central nervous system (CNS) (Boyke et al., 2008; Dayan and Cohen, 2011; Draganski and May, 2008; Hüfner et al., 2011; Scholz et al., 2009; Taubert et al., 2016). Studies using magnetic resonance (MR) imaging have provided evidence for grey (GM) and white matter (WM) changes accompanying the acquisition of specific skills in brain regions engaged by the task or activity (Draganski and May, 2008; Zatorre et al., 2012). Volumetric GM changes have been observed in the hippocampus after acquiring expertise in dancing (Hüfner et al., 2011) and spatial navigation tasks (Maguire et al., 2000) and in the putamen of skilled pianists (Granert et al., 2011). Even when skills are acquired by non-experts, over much shorter timescales, GM changes are apparent in the motor cortex (M1) after one hour of balance training (Taubert et al., 2016) or after 16 days of temporary contralateral limb immobilisation (Langer et al., 2012), in the left parahippocampal and right superior temporal gyrus after three weeks of unicycling (Weber et al., 2019) and in the left hippocampus and bilateral nucleus accumbens after 3 months of juggling (Boyke et al., 2008). The trajectory of these changes suggests that learning-related changes follow a sequence of expansion, selection, and renormalization, although the precise nature of these neural processes remains unresolved (Boyke et al., 2008; Draganski et al., 2006; Fu and Zuo, 2011; Makino et al., 2016; Scholz et al., 2009; Weber et al., 2019; Wenger et al., 2016).

Training-induced WM volume and microstructural changes—inferred indirectly from diffusion tensor imaging (DTI)—have been documented in parietal (Lakhani et al., 2016; Taubert et al., 2011), fronto-parietal^16^ and temporo-occipital regions (Deng et al., 2018) following sustained motor practice. Both GM and WM changes on MR neuroimaging have been associated with visual, somatosensory and motor cortical neuroplasticity, performance improvements, oligodendrogenesis and myelination in mice (Badea et al., 2019; Gibson et al., 2014; Lamprecht and LeDoux, 2004; Theodosis et al., 2008). More recently, MR techniques utilising advanced computational modelling of multi-parameter mapping (MPM) have been developed (Dick et al., 2012; Freund et al., 2019; Helms et al., 2008; Leutritz et al., 2020; Natu et al., 2019; Sereno et al., 2012; Weiskopf, 2013), allowing detailed elucidation of microstructure including myelin and iron (Natu et al., 2019), which have been used to demonstrate changes indicative of demyelination in the CNS of humans following spinal cord injury (David et al., 2019; Freund et al., 2019).

Hitherto, the motor tasks used to explore training effects have limited applicability in rehabilitation, with unicycling and juggling of little relevance to most patients with significant neurological injury or disease. We aimed to study the evolution of GM and WM volume changes—uniquely adding longitudinal, myelin-sensitive MPM analyses—in the brains of healthy individuals during 1 month of intensive sensorimotor training in a challenging, yet feasible, task for neurological patients and healthy individuals alike, and which was applicable to both the upper and/or lower limbs (Prahm et al., 2017). Participants learned to perform increasingly complex and rapid sequences of movements of either their upper or lower limbs in response to visual and auditory cues. Using serial MR imaging before, during and after training, we aimed to characterise (i) the spatial and temporal evolution of training-induced changes across the primary motor system and hippocampus, (ii) the somatotopic changes specific to upper or lower limb training and (iii) associations between structural changes and improvements in task performance.

The training task selected for use in this study is based on an open source dance and rhythm game used for recreation and rehabilitation, known for its modifiability, variable difficulty and for its capacity to provide a challenging, fun and motivating game experience (Prahm et al., 2017). Playing is known to recruit key attentional, integrative and sensorimotor brain networks (Eggenberger et al., 2016; Noah et al., 2015) and is expected to engage hippocampal circuits involved in the consolidation of motor sequence learning (Albouy et al., 2008; Long et al., 2018), with the combination of multijoint physical activity and cognitive and sensorimotor stimulation particularly likely to promote consolidated learning (Kempermann et al., 2010; Rehfeld et al., 2017). While activation of some neural elements will be common, irrespective of the limb trained, we anticipated differences between those training with the upper and lower limbs, in regions with well-established somatotopy, including the sensorimotor and corticospinal systems.

## Materials and Methods

### Participants

Thirty-two healthy, right-handed adult (age range: 18-50 years) males were recruited. All had normal or corrected-to-normal vision, no history of psychological or neurological disease, and no contraindications to MR. All participants were naive to the experimental setup. The first six participants were block randomised to either upper, lower or no training groups, after which an age-matching algorithm was used to ensure comparability to our typical spinal cord injury patients (young to middle-aged males) for future studies including patient cohorts.

### Standard protocol approvals, registrations, and patient consents

The study was carried out in accordance with Good Clinical Practice and the Declaration of Helsinki and approved by the Zurich cantonal ethics committee (KEK-2013-0559). All participants gave written, informed consent.

### Training task

Participants undertook training over four consecutive weeks (60 minutes training, 4x per week, figure 1A). They were not allowed to train on the task between Day 28 and Day 84, when an assessment of performance retention was made. Upper and lower limb training utilised StepMania 5 Beta 3 (www.stepmania.com) for Windows 7 (Microsoft, La Jolla, CA) and an input device dependent on which limbs were to be trained (see below). During the game, arrows matching symbols on the input device (← ↑ → ↓) scrolled up the screen while a popular song played from the speakers (see accompanying video). The participant was tasked with selecting and activating the correct symbol at the precise moment the scrolling arrow overlapped with a set of static arrows at the top of the screen with the moment of overlap synchronised to the beat of the music using the Dancing Monkeys script (Karl O’Keeffe, https://monket.net/dancing-monkeys/). The script generates patterns of arrows of varying difficulty but excludes sequences that would be impossible to respond to. Each bout lasted 120 seconds, after which the participant was given immediate visual feedback in the form of a percentage score (accurate response within 45ms: 2pts, 45-90ms: 1pt, >90ms: no score; cumulative score expressed as a % of maximum possible points). As the pattern of arrows differed for each bout, participants achieved improvement not by rote learning of a series of movements but by developing optimal strategies for adapting to the varying patterns (Orrell et al., 2006), however, optimal response involves identifying and executing multi-step responses to frequently-encountered patterns of arrows as they are revealed. Each training session comprised 15 bouts. Progress in the training involved moving through increasingly difficult levels whereby the number, pattern complexity and scroll speed of the arrows increased. The next level was unlocked when three non-consecutive scores of ≥ 80% were achieved within a level. Demotion to the previous level was mandated by three consecutive scores of ≤ 30%.

**Fig. 1:**
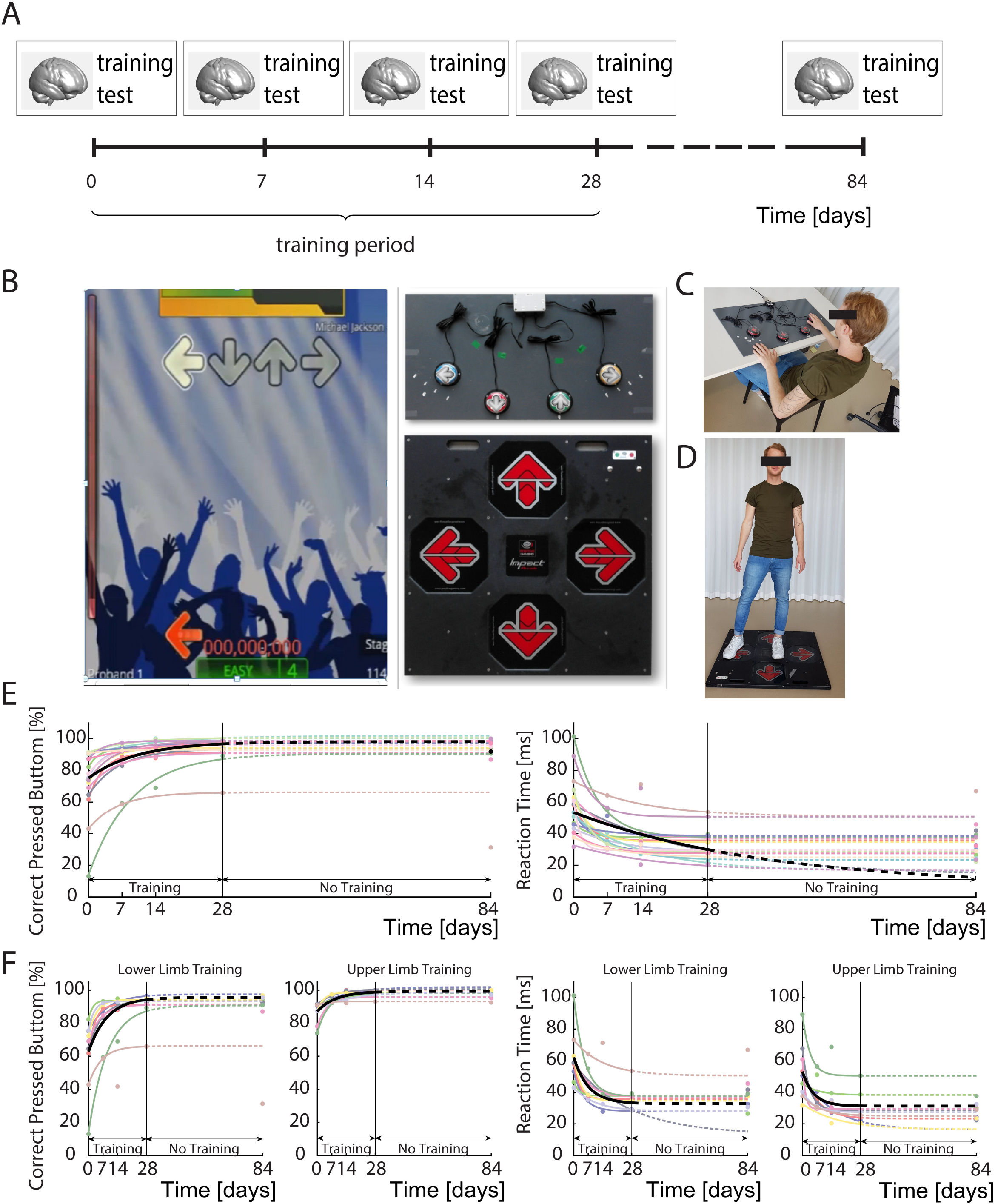
Experimental design, training task and behavioural data (see accompanying video). The experimental design (A) included MRI acquisition and training assessments at baseline (day 0) during the training period (days 7, 14 and 28) and at final retention assessment (day 84). Sixty minutes of supervised training in a motor-skill task was undertaken four times per week for 4 consecutive weeks (B) whereby participants were required to activate inputs with their hands or feet (depending on whether they were allocated to the upper (C) or lower limb (D) training groups) in response to aural rhythmic and visual stimuli in the dance game Stepmania. The participant was tasked with selecting and activating the correct symbol at the precise moment the scrolling arrow overlapped with a set of static arrows at the top of the screen. Behavioral improvement, defined as the percentage of correct stimulus responses (%CSR), and response time (RT; the deviation in ms from the ideal response) were measured during a formal, standardized performance test at weekly intervals (see methods). Individual values for these metrics are plotted as coloured dots (E and F), while participant-specific behavioral curves defined by the function e (*y* = *α* − *β e^−γ*,t^*) (see methods), were computed (coloured lines) along with the corresponding group mean (black line). The dashed lines connect the last training point (Day 28) with the retention test on Day 84. All participants demonstrated improvement in the task over time (p<0.05) and that %CSR evolution was significantly different (p=0.004) between the upper and lower limbed trained groups, while RT was not.

Participants allocated to the lower limb training group used a dance platform (Impact Dance Platform, Positive Gaming BV, Haarlem, Netherlands) as the input device and effectively learned to “dance” to the songs (figure 1 B, D). The platform was placed on the floor in a large open space facing a TV monitor on which the game was displayed. Those allocated to the upper limb training group sat facing a table on which a custom-made platform, designed to be analogous to the lower limb platform, was placed (figure 1 B, C). Participants were instructed to use their left hand for ← and ↑ and their right for → and ↓. A laptop was used to present the game.

Each participant progressed through the levels at different paces and were consistently challenged. To assess changes in task performance in a standardised manner, participants were assessed immediately prior to training (baseline) and again on days 7, 14, 28 of training and finally on day 84. These performance tests were conducted using a pre-programmed sequence of arrows including segments at 60, 80, 100 and 120 bpm and featuring patterns of increasing complexity for total of 3min 20sec set on a neutral, black background with the music replaced with a metronome.

Importantly, the sequence of arrows presented differed for each test. Thus, each test was controlled in terms of overall difficulty but learning of the specific sequences was not possible. Cues (← ↑ → ↓) were balanced within and across segments to ensure no laterality effect. Response and response time data in these standardised performance tests were used as the basis for the behavioural analyses.

### Behavioural analyses

Behavioural data were analysed using bespoke Matlab 2016b (The MathWorks, Natick, MA, USA) routines. Performance was measured as the percentage of correct stimulus responses (%CSR; defined as accurate inputs within 90ms of a cue); and response time (RT), defined as the mean absolute delay in ms between cue overlap and correct inputs. These behavioural measurements were computed for each subject and each side (left/right). In order to evaluate the improvement of each subject, subject-specific model parameters were estimated as follows.

An exponential model of both behavioural parameters was computed over the training period using the following nonlinear regression model:

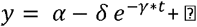

where y is the training parameter (%CSR or RT); t is the training timepoint (baseline, day 7, 14 and 28), with random error ⍰, assumed to be normally distributed with a mean of 0. *α* is the ‘asymptote’, i.e. the value the training parameter eventually settles on over time (the plateau); *δ* is the ‘acquisition climb’, the extent of improvement from baseline to asymptote; and *γ* indicates the shape of the trajectory (i.e., how quickly it converges to asymptotic performance). These parameters were estimated for each participant by fitting the model to the subject’s specific behavioural parameters. Parameters *δ* and *γ* were used to investigate: (i) the improvement of all participants; (ii) if there are differences in training upper and lower limbs; (iii) if there are differences in training left and right side; and (iv) if any differences are associated with MRI changes. The first three questions were assessed with Stata 15.0 (Stata Corp, College Station, TX); while the latter was assessed with SPM (Statistical Parametric Mapping version 12; https://www.fil.ion.ucl.ac.uk/spm). The distribution of performance metrics (in terms of δ and *γ*) was first assessed (using the Shapiro-Wilk test for normality) and analysed for all trained participants using one-sample Wilcoxon tests. To assess performance differences when training the upper versus lower limb, a two-sample Wilcoxon test was performed on the *γ* and *β* of all trained participants. Finally, to assess if the acquired skill was preserved at follow-up, a Wilcoxon matched-pairs signed-rank test was performed on %CSR and RT at days 28 and 84.

### MRI acquisition

MRI data were acquired on a single 3T Siemens Skyra^flt^ scanner before training (baseline), during the training at 7, 14, 28 days, and at 84 days follow-up. If scanning and training were scheduled on the same day, the training took place after the scan.

All participants underwent a multi-parametric mapping (MPM) MRI protocol to furnish estimates of longitudinal relaxation rate (R1=1/T1), magnetization transfer saturation (MT) and transverse relaxation rate (R2*)(Helms and Dechent, 2009; Leutritz et al., 2020; Weiskopf et al., 2013, 2011). The MPM protocol acquires three volumes using a 3D multi-echo sequence (based on the Siemens “gre” FLASH product sequence) with 1 mm isotropic resolution, field of view (FoV) = 240×256×176 mm^3^. Total acqusition time was 23 mins and used parallel imaging with an acceleration factor of 2 in each phase-encoding direction and subsequent reconstruction with the generalized auto-calibration partially parallel acquisition algorithm (GRAPPA). The readout bandwidth was 480 Hz/pixel. Each set of echoes was acquired using a different TR and radio-frequency (RF) excitation flip angle (α) to achieve images with either T1-weighting: 25 ms / 23°, proton density (PD)-weighting: 25 ms / 4°, or MT-weighting: 37 ms / 9°, with an off-resonance MT saturation pulse applied prior to excitation. Echoes were acquired at 6 equidistant echo times (TE) from 2.46 ms to 14.76 ms for all weightings, with an additional 2 echoes at 17.22 and 19.68 ms for the PD-weighted and T1-weighted volumes.

### MRI processing

A total of five MRI scans were acquired for each participant. MPM maps were generated, using UNICORT to correct transmit field inhomogeneities, with the hMRI toolbox (Tabelow et al., 2019) in SPM12 (Wellcome Trust Centre for Neuroimaging, London, UK, http://www.fil.ion.ucl.ac.uk/spm) and Matlab. All maps were pre-processed using an analysis pipeline for longitudinal data (Ziegler et al., 2018), in which MT maps were first skull-stripped and then used for longitudinal registration within-participants. This step was based on a generative model in which each image volume was registered to a subject-specific average map, combining non-linear and rigid-body registration with corrections for intensity bias artefacts. This procedure generated participant-specific, midpoint maps with corresponding deformation fields.

Second, a unified segmentation was applied to the subject’s midpoint map generating probability maps of GM, WM, and cerebrospinal fluid (CSF). Third, nonlinear template generation and image registration was applied to subject-specific, midpoint GM and WM tissue maps—and the template was registered to Montreal Neurological Institute (MNI) space using an affine transform based on Dartel (Ashburner, 2007). Fourth, all MPM maps were warped to MNI space using transformations obtained in previous steps. Finally, all MPM maps in MNI space were spatially smoothed using a 5-mm (in the GM) and 3-mm (in the WM) full-width at half-maximum (FWHM) in order to minimize partial volume effects of GM/WM tissue by appropriate tissue segment weighting (Draganski et al., 2011).

To relate the quantitative MPM metrics to conventional morphometric approaches (such as voxelbased morphometry [VBM]), MT maps were additionally pre-processed using the longitudinal VBM analysis from CAT12 toolbox (http://www.neuro.uni-jena.de/cat/). Specifically, this longitudinal VBM analysis includes, first the realignment of all time points for each subject using an inverse-consistent rigid registration, which includes bias-correction; then, the realigned MT were segmented into GM, WM and CSF. Next, deformation fields to the MNI space, were computed using a non-linear spatial registration (Dartel) and averaged (Ashburner, 2007). The resulting mean deformations were applied to the tissue segmentations (GM and WM probability maps) of all time points, modulated with the Jacobian determinant of the deformation; and spatially smoothed using isotropic Gaussian smoothing of 6-mm (in the GM) and 4-mm (in the WM) FWHM. For statistical analysis, we excluded all voxels with a GM probability below 0.2 and WM probability below 0.6 to minimize contribution of partial volume effects near GM/WM borders.

### MRI Statistics

#### Training-induced structural changes

We used the SPM sandwich estimator (SwE) toolbox (http://www.nisox.org/Software/SwE) to assess: (i) training-induced brain changes; (ii) topological changes and (iii) how training improvements (in terms of δ and *γ* from the behavioural analysis) were related to structural trajectories. The SwE was developed for efficient, voxel-based longitudinal image analysis and it is based on a marginal model where the expected variability is described as a function of predictors in a design matrix, while additionally accounting for correlations due to repeated measurements and unexplained variations across individuals as random effects. The model incorporated a linear (since baseline) and a quadratic time factor to model persistent (linear term) or transient (quadratic term) training-induced brain changes defined as the difference between the combined trained (upper + lower limb training groups) with the untrained group. The SwE model is similar to conventional mixed effects general linear model (GLM), specified in terms of a design matrix containing subject-level predictors (such as time, time^2^ and covariates). The parameters of this model are estimated in the usual way for every voxel:

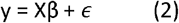

Here, y denotes the tissue volumes or quantitative MPM metrics at multiple timepoints during the training period, X denotes the design matrix including predictors [intercept, time, time^2^], group interactions and covariates [age, TIV] and *ϵ* indicates random effects with between-subject variance components with mean 0 (for details see (Guillaume et al., 2014)). The time and time^2^ parameters were mean-centred and orthogonalized which enabled the identification of both linear and concave/convex, u-shaped trajectories. The same model was used to assess somatotopic effects comparing lower-limb with upper-limb trainees.

For the analysis of the association between MRI and motor learning parameters, we enriched the models by adding the learning parameters sequentially. In particular, we investigated the association between linear and quadratic structural changes with behavioural improvement (δ), speed of improvement (*γ*) and baseline value (α-δ) for each participant-specific model, using both training parameters (%CSR and RT) as well as their interaction with change over time and time^2^. These variates allowed us to assess how linear or quadratic brain changes were associated with training improvements. In all models, the mean age of a participant and total intracranial volume (TIV) were included as covariates of no interest. All results with p-values of 0.05 (FDR corrected) were considered significant. In order to preclude potential, non-specific global compounds (e.g. due to residual biases after UNICORT correction), WM changes in the frontal pole of the control group were assessed: no changes were observed (p > 0.05, FDR corrected). To assess whether structural changes persisted after a period of no training, a paired t-test (p<0.05 family-wise error at the cluster level) was performed comparing scan data at follow-up (day 84) and end of training (day 28).

#### Regions of interest (ROI)

Regions of interest (ROI) were used to summarise training-related changes in structure for subsequent tests of coherent changes between ROI. The ROI were chosen based on prior studies reporting GM and WM changes in response to upper and lower limb training (Boyke et al., 2008; Draganski et al., 2006; Hüfner et al., 2011; Lakhani et al., 2016; Schlegel et al., 2012; Scholz et al., 2009; Wenger et al., 2016) to include NS networks likely to be recruited by the various elements of the task; including consolidation of motor sequence learning (Long et al., 2018). These comprised the primary sensory area (S1) and M1, cranial corticospinal tract (CST) and the hippocampus-entorhinal cortex (EC) formation. Hippocampal and CST sub-regions were defined based on Oxford Centre for Functional MRI of the Brain Software Library (FSL) Templates in MNI space; while S1 and M1 were defined using the anatomy toolbox in SPM (Eickhoff et al., 2007, 2005).

#### Coherent changes in microanatomy of the corticomotor system and within the hippocampus

To investigate coherent (i.e., correlated but time-lagged) changes in structural metrics within and between the corticomotor system and hippocampus, the mean value of structural metrics from the significant clusters in the trained group was extracted from the maps in MNI space for all participants. A time-lagged version of each cluster was computed by shifting the group mean values for each session by one-time step in the training period (baseline, days 7, 14 and 21). Finally, time-lagged changes in the corticomotor system and hippocampus were tested using a mixed effects GLM. In other words, we looked for correlations between changes in one cluster (i.e. mean R1 from the left M1) with time-lagged changes in another (i.e. mean R1 from left cerebral peduncle cluster at the preceding timepoint). All results with p < 0.05 (uncorrected) were considered significant.

## Results

### Demographics and behavioural results

The three training groups did not differ with respect to age (upper limb training [n=9, age=34±10.22 SD years]; lower limb training [n=9, age=38.67±13.22 SD years]; no training [n=14, age=38.71±10.98 SD years; ANOVA: F=0.49, p=0.61]). All trained participants improved (in terms of δ) in both %CSR (p<0.001, z= 3.72) and RT (p<0.001, z=3.72), over the 28 days of training (figure 1E). At baseline, the combined group of trained participants group achieved 76.75 %CSR (interquartile range [IQR]: 69.11-90.38 %) improving by a median of 18% (IQR: 11–23%) by 18 days (time = 3/*γ* for 95% improvement, with *γ* = 0.17, IQR: 0.11-0.26). Median RT was 55.8 ms (IQR: 45.71-63.04 ms) at baseline, and trained participants improved by a median 25.77 ms (IQR=15.00 – 34.82) reaching a plateau at 18 days (time = 3/*γ* for 95% improvement, with *γ* = 0.17, IQR: 0.10-0.28).

At baseline, lower limb trainees achieved 69.11 % CSR (IQR: 61.78-75.16) and RT was 58.83 ms (IQR: 53.35-63.04); while upper limb trainees achieved 90.38 % CSR (IQR: 87.73-90.76 %) with an RT of 50.57 ms (IQR: 40.57-60.93 ms). This allowed participants training the lower limbs to improve more in terms of % CSR (22% [IQR 22-29] versus 11% [9-12]) compared to upper-limb trainees (p= 0.004, two-tailed test), while improvement in RT was similar between the groups (lower limb: 30.05 ms, [IQR 13.−8 - 34.82] versus upper limb: 25.4 ms, [22.58-28.4], p= 0.895, two-tailed test).

Specific improvements in %CSR and RT for inputs exclusively delivered by the left or right side were not significantly different (p= 0.05, two-tailed test). Between Day 28 and 84, no significant differences between % CSR and RT were observed (p=0.930 and p=0.349 respectively; figure 1F).

### Training-induced structural changes common to both upper and lower limb trainees

#### Corticospinal system

Across the training period (Day 0 – 28), in all trainees compared to the untrained group, transient R1 and R2* decreases following a convex, u-shaped course were observed in the left primary sensorimotor cortex (R1: z=2.82, p=0.048 FDR corrected and z=3.099, p=0.047 FDR corrected; R2*: z=4.02, p=0.01 FDR corrected) and a trend towards the same in the right primary sensorimotor cortex for R2* (z=3.31, p=0.051). R2* also showed a trend towards a linear decrease in a region in the left primary sensorimotor cortex (z=2.93, p=0.052). In the cranial CST at the level of the cerebral peduncle, transient R1 decreases following a convex, u-shaped course were observed (z=3.502,p=0.021 FDR corrected, z=3.199,p=0.025 FDR corrected, left and right respectively), and at the level of the right brainstem (z=3.607 p=0.012 FDR corrected; z=2.944 p=0.033 FDR corrected, figure 2, table 1), while a similar association, observed in the CST at the level of the right posterior limb of the internal capsule, did not reach statistical significance (z=1.938, p=0.052 FDR corrected).

**Fig. 2:**
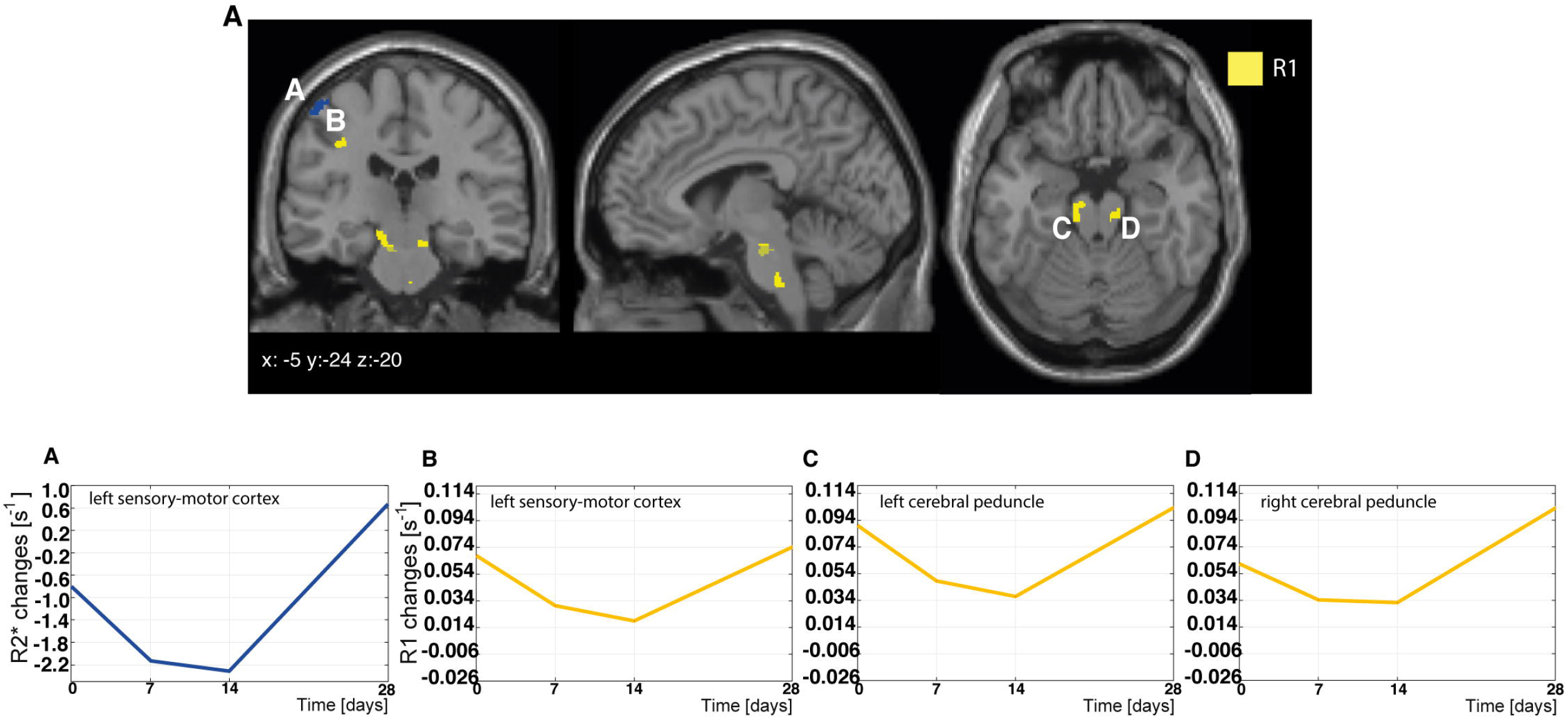
Training-induced changes in the corticomotor system during training. Transient, quadratic microstructural changes (R1 (yellow), R2* (blue)) were observed in the bilateral corticospinal tract (i.e. cerebral peduncle) and left motor cortex. All observed changes followed a convex trajectory as shown in the longitudinal plots in B, C and D.

**Table 1:** Significant differences in linear/quadratic changes during day 0-28 between the combined trained vs. untrained group. SPM longitudinal SwE results table testing for linear/quadratic differences between trained and untrained group on grey matter (GM) volume, myelin-sensitive R1 and MT maps. We report p-values from non-parametric voxelwise FDR (p<0.05). Non-significant trends of interest in italics.

| ROI | Map | Contrast | p value | z value | x<br>[mm] | y<br>[mm] | z<br>[mm] |
| --- | --- | --- | --- | --- | --- | --- | --- |
| left cerebral peduncle | R1 | quadr Trained>NotTrained | 0.021 | 3.502 | -15 | -24 | -15 |
| right cerebral peduncle | R1 | quadr Trained>NotTrained | 0.025 | 3.199 | 17 | -12 | -8 |
| right posterior limb of the internal capsule | R1 | quadr Trained>NotTrained | 0.052 | 2.938 | 20 | -15 | -5 |
| left CST at the level of the brainstem | R1 | quadr Trained>NotTrained | 0.012 | 3.607 | -5 | -18 | -26 |
| right CST at the level of the brainstem | R1 | quadr Trained>NotTrained | 0.033 | 2.944 | 3 | -27 | -42 |
| left primary motor cortex | R1 | quadr Trained>NotTrained | 0.047 | 3.099 | -33 | -20 | 44 |
| left primary motor cortex | R2* | quadr Trained>NotTrained | 0.010 | 4.017 | -50 | -21 | 57 |
| *left primary motor cortex* | *R2\** | *linear Trained<NotTrained* | *0.052* | *2.932* | *-29* | *-35* | *51* |
| *right primary motor cortex* | *R2\** | *quadr Trained>NotTrained* | *0.051* | *3.307* | *33* | *-44* | *69* |
| left primary sensory cortex | R1 | quadr Trained>NotTrained | 0.048 | 2.824 | -38 | -24 | 39 |
| **Hippocampus** |  |  |  |  |  |  |  |
| left entorhinal cortex | MT | linear Trained>NotTrained | 0.020 | 3.790 | -24 | -9 | -38 |
| left entorhinal cortex | R1 | linear Trained>NotTrained | 0.047 | 3.099 | -27 | -11 | -41 |
| right entorhinal cortex | R1 | linear Trained>NotTrained | 0.039 | 3.038 | 21 | 2 | -30 |
| left anterior hippocampus | Volume (GM) | quadr Trained<NotTrained | 0.036 | 3.036 | -24 | -17 | -21 |
| left anterior hippocampus | MT | quadr Trained>NotTrained | 0.034 | 2.766 | -20 | -17 | -18 |
| left dorsal hippocampus | R1 | quadr Trained>NotTrained | 0.040 | 3.232 | -33 | -36 | -9 |

#### Hippocampus

In the hippocampal formation, a transient R1 decrease in the dorsal hippocampus (z=3.232, p=0.040 FDR corrected) and MT decrease in the anterior hippocampus (z=2.766, p=0.034 FDR corrected), were observed in trainees but not in untrained controls, again following a convex pattern. These transient decreases in R1 and MT were accompanied by a similarly transient volumetric increase in the left anterior hippocampus (z=3.036, p=0.036 FDR corrected; Figure 3 and Table 1). A linear increase in R1 over time was observed in the bilateral EC (z>3, p< 0.05 FDR corrected), while MT linearly increased in the left EC (z=3.79, p= 0.02 FDR corrected) in trainees compared to those who were untrained (Figure 3 and Table 1). No significant changes in macro- or microstructure were observed between days 28 and 84 in any ROIs.

**Fig. 3:**
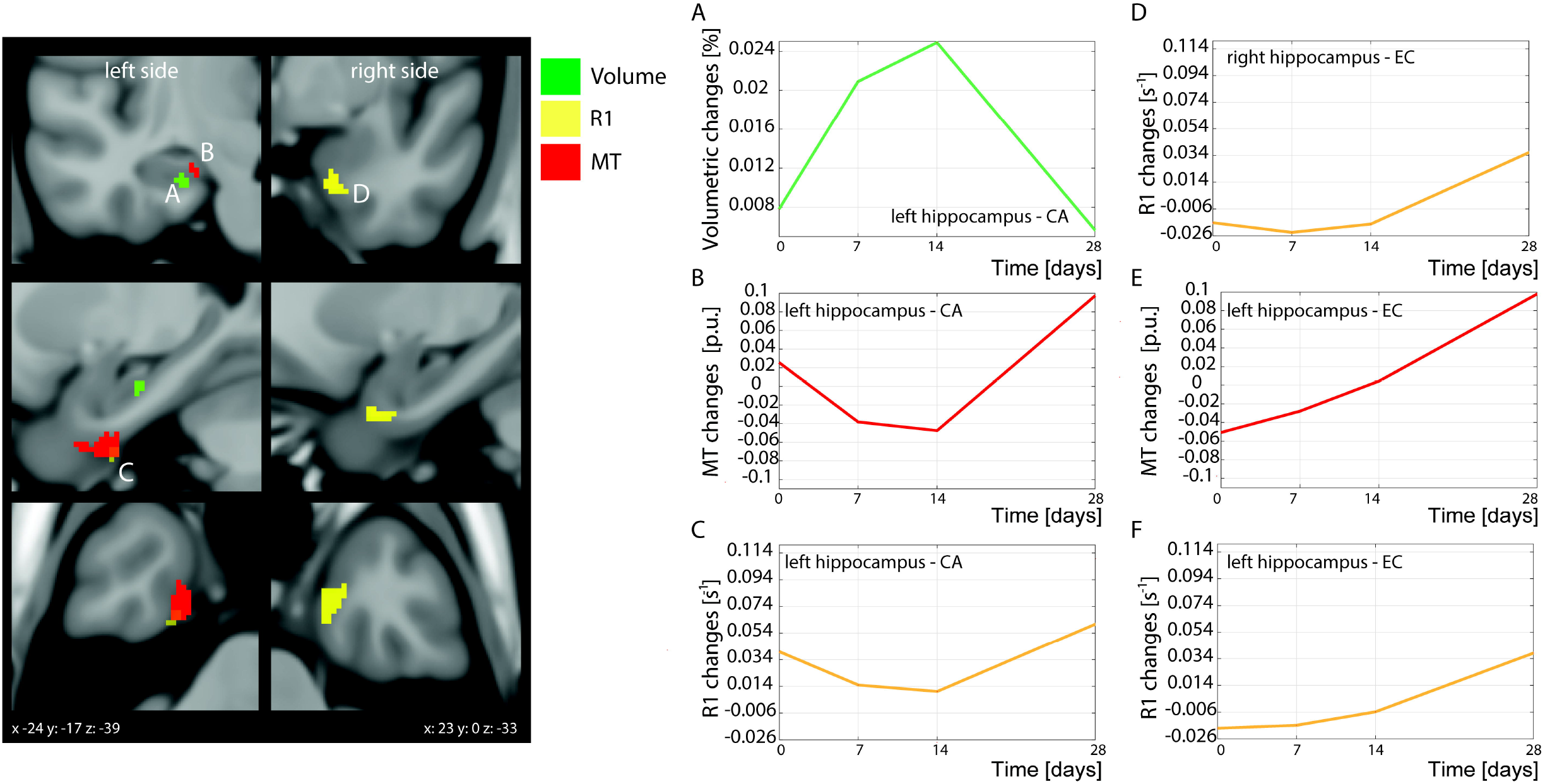
Training-induced changes in the hippocampus. Transient, quadratic grey matter volume changes where observed in the left hippocampal cornu ammonis (CA) following a concave course, (A) while microstructure (MT and R1 respectively shown in B and C) followed a convex, u-shaped course. Linear increases were observed in the bilateral entorhinal cortex (EC) for R1 (D and F) and left hippocampus-EC for MT (E).

### Somatotopy of lower vs. upper limb training

When comparing upper and lower limb trainees, we found a larger, transient (i.e. quadratic) MT change in the lower-limb area of the CST at the level of the left internal capsule (z=3.0, x=-24, y=-20, z=9, p=0.023, FDR corrected), and left cerebral peduncle (z=3.079,x=-21, y=-21, z=-8, p=0.047 FDR corrected; figure 5) in lower limb trainees.

### Associations with performance improvements

Improvements in correct stimulus response (%CSR) were associated with quadratic volumetric changes in the left anterior hippocampus (z=2.885, p=0.019 FDR corrected, table 3), indicating that larger transient volume increases (represented by a negative quadratic term) predict greater improvement in %CSR. A positive association between these volumetric changes with improvement in RT (z=2.834, p=0.028 FDR corrected; table 3) was seen in the same region, suggesting that larger transient volumetric increases are related to shorter RT. Positive associations were found between linear MT changes in the right EC, and more rapid improvement (*γ*) in % CSR and RT (z=3.568, p=0.04 FDR corrected; z=3.797, p=0.028 FDR corrected, respectively; figure 6 and table 3). A positive association between R2* changes with improvement in RT (z=4.0, p=0.022 FDR corrected; table 3) was seen in the left primary sensorimotor cortex, suggesting that larger linear R2* increases are related to shorter RT. Baseline %CSR correlated negatively with quadratic MT changes in the left EC (z=3.607, p=0.026 FDR corrected; table 3).

### Coherent plasticity in the motor cortex, the corticospinal tract and the hippocampus

At any given timepoint, there was a significant positive correlation between changes in R1 in the left M1 and changes in R1 in the left CST at the level of the left cerebral peduncle at the preceding timepoint (coefficient= 0.138, p=0.026, CI:0.016 to 0.258). Similarly, there was a significant negative correlation between R1 changes in the left EC and R1 changes in the left M1 at the previous timepoint (coefficient= −0.360, p=0.003, CI: −0.598 to −0.122), with R1 decreases in M1 preceding R1 increases in the left EC. A further significant negative correlation was observed between R1 changes in the left EC and R1 changes in left dorsal hippocampus at the previous timepoint (coefficient=-0.298, p=0.006, CI: −0.509 to −0.086), with decreases in R1 in left dorsal hippocampus preceding R1 increases in the left EC (figure 4 and table 2).

**Fig. 4:**
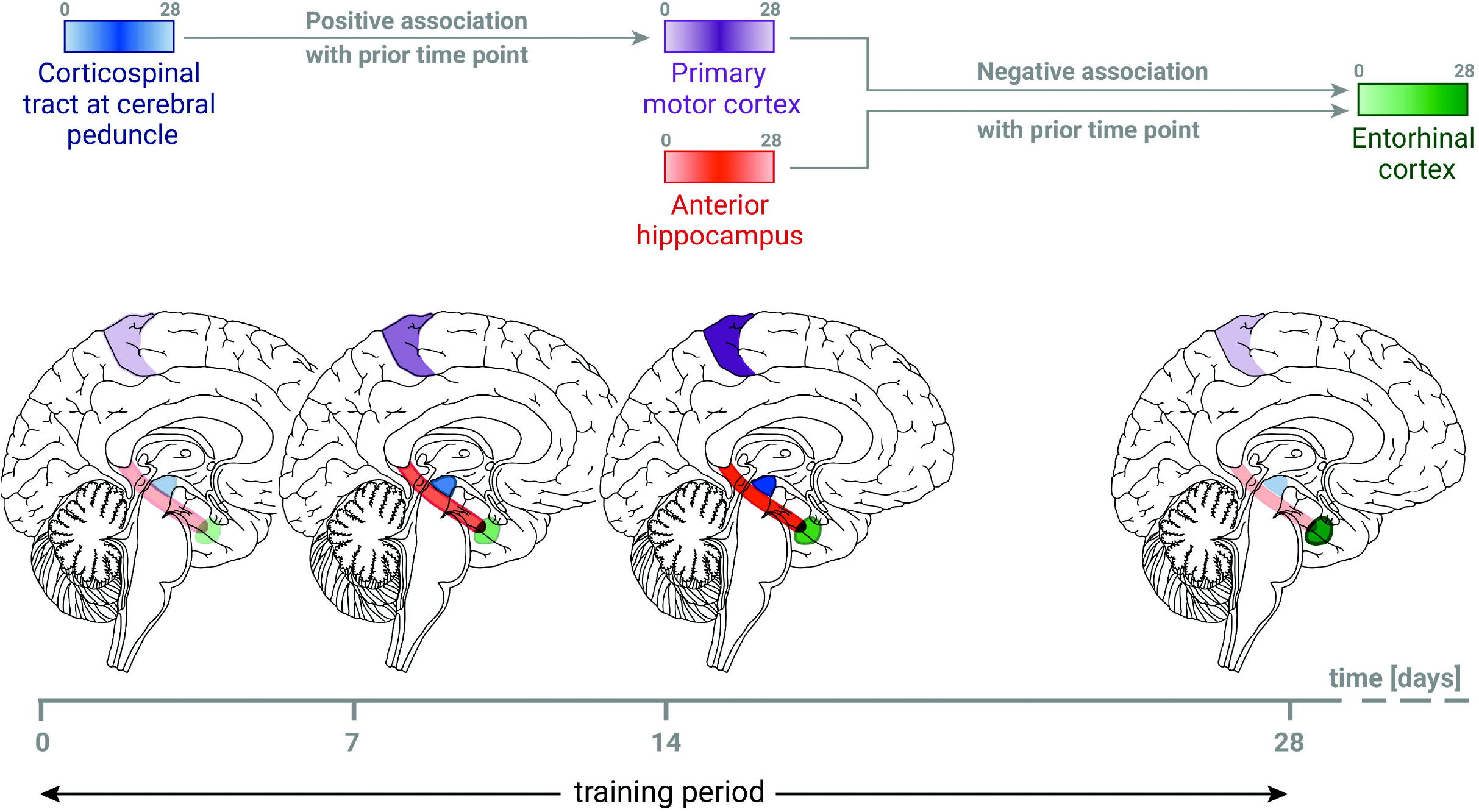
Schematic representation of the coherent R1 changes between corticomotor and hippocampi circuits over time during training. A positive correlation was observed with R1 increases in the left corticospinal tract (blue) at the level of the cerebral peduncle preceding later R1 increases in left primary motor cortex (purple). Elsewhere, negative temporal correlations were observed, with increases in R1 in the left primary motor cortex or the left anterior hippocampus (red) associated with R1 decreases in the left entorhinal cortex (green). Note, the gradient colour changes within the boxes and brain illustrate the non-linear changes observed in the corticospinal tract, primary motor cortex and anterior hippocampus while linear changes were observed in the entorhinal cortex.

**Table 2:** Time lag analysis between myelin-sensitive R1 changes along the corticomotor system and with and within the hippocampus.

| **R1 map temporal association** | **Coefficient** | **Standard deviation error** | **z value** | **p value** | **95% Confidence interval** |
| --- | --- | --- | --- | --- | --- |
| Left primary motor cortex and left cerebral peduncle one timepoint prior | 0.137 | 0.062 | 2.220 | 0.026 | 0.016 to 0.258 |
| Left hippocampus (entorhinal cortex) and left primary motor cortex one timepoint prior | -0.360 | 0.121 | -2.970 | 0.003 | -0.598 to -0.122 |
| Left entorhinal cortex and left dorsal hippocampus one timepoint prior | -0.298 | 0.108 | -2.760 | 0.006 | -0.509 to -0.864 |

**Table 3:**
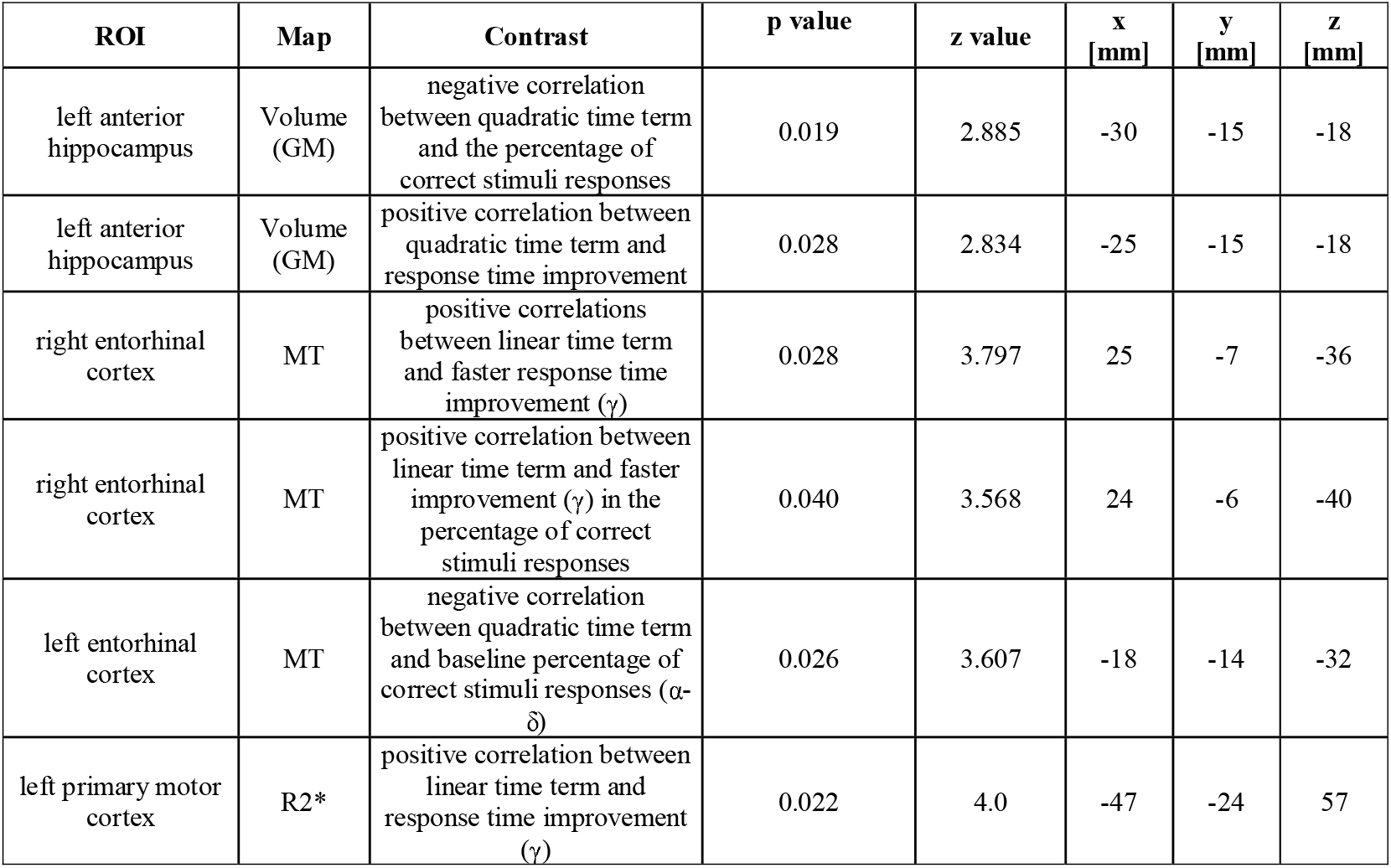
Correlation table. SPM longitudinal SwE results table testing for correlations between linear/quadratic structural changes and behavioural improvements. We report p-values from non-parametric voxel-wise FDR (p<0.05).

## Discussion

This study provides novel insights into the character of brain changes accompanying the acquisition of skill in the motor learning game Stepmania. Among trainees, temporal and spatial microstructural changes in the WM and GM of the primary motor system revealed an intricate interplay with both transient and persisting hippocampal changes, the magnitude of which was associated with improved performance. Upper versus lower limb somatotopy was reflected in differences in myelin-sensitive MT at the level of the left internal capsule and cerebral peduncle. These findings speak to putative mechanisms of neuroplasticity that underwrite motor learning—with potentially transferable insights for motor rehabilitation and clinical practice.

### Training-induced structural changes

#### The corticospinal system

R1 and MT are increasingly accepted as surrogate markers of tissue myelin content (Natu et al., 2019; Schmierer et al., 2004) while R2* is known to be highly sensitive to iron deposition (Ghadery et al., 2015; Langkammer et al., 2010). Over 1 month of active training, we observed transient (i.e. following a convex course) R1 and R2* decreases and subsequent increases in the sensorimotor cortex and, for R1, in the bilateral CSTs of trainees. Such dynamic changes in R1 and R2* may represent a relatively reduced proportion of myelin due to training-induced angiogenesis or gliogenesis (Badea et al., 2019; Fields, 2015) and are in keeping with the early literature on local tissue volume expansion in response to training (Draganski and May, 2008; Zatorre et al., 2012). Such tissue expansion may also lead to changes in iron consumption as active myelination is taking place (Sampaio-Baptista et al., 2020, 2013). Transient vasculature changes have been demonstrated *in vivo* where physical exercise induced a temporary, histologically-confirmed increase in vascular volume in the adult simian M1 (Rhyu et al., 2010), although human neuroimaging studies analysing cerebral blood volume did not corroborate this (Thomas et al., 2016). Oligodendrogenesis has been observed in recruited brain regions in trained rodents (Gibson et al., 2014) and the inactivation of oligodendrocyte precursor cells (OPC) leads to impaired skill learning (McKenzie et al., 2014). However, in adult humans, the oligodendrocyte population is stable with a low turnover rate, suggesting myelination in response to training is instead carried out by mature oligodendrocytes, rather than differentiation from OPCs (Yeung et al., 2014). As both angiogenesis and gliogenesis are based on changes in non-neural substrates common to both GM and WM, either or both could underlie the transient changes observed. Although participants trained both left and right limbs equally, the transient changes observed in M1 were only observed on the left side. It is possible that our group of exclusive right-handers may have experienced greater skill improvement in their dominant limb, with performance plateau driven by poorer performance of the left limb, leading to more pronounced changes in the left brain (McGrath and Kantak, 2016). Alternatively, the well-established relative proclivity of the left hippocampus to undergo plastic change (Boyke et al., 2008; Weber et al., 2019) may, through the cortico-limbic interplay revealed in this study and detailed later, lead to more pronounced microstructural changes in left hemisphere regions over the timescales assessed here.

Somatotopic analysis revealed greater transient increases in MT contrast (figure 5) in the left internal capsule and lateral left cerebral peduncle of lower-limb trainees compared to those who trained the upper limb. This is in keeping with somatotopic anatomy of the lower limb corticospinal tract and the fact that lower limb trainees began from a lower performance level at baseline, with greater absolute improvement (Δ% CSR) achieved by this group. These findings are supported by animal studies in which greater task improvement was correlated with increased FA following learning (Sampaio-Baptista et al., 2013), with improvement related to activity dependent myelination, which selectively increased conduction velocity through white matter circuits (Fields, 2015; Hofstetter et al., 2013). The internal capsule and cerebral peduncle represent particularly invariable elements of the human intracranial CST (Reich et al., 2006; Song, 2009), rendering them potentially important loci for the study of microstructural learning effects.

**Fig. 5:**
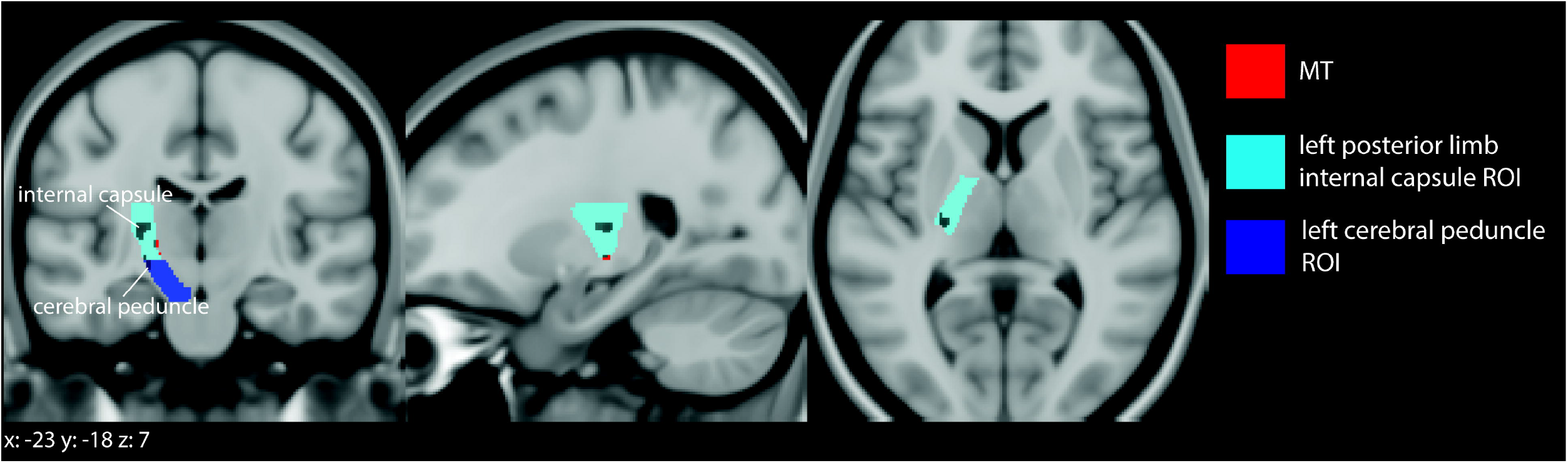
Somatotopic differences associated with training the upper vs lower limb. Training of the lower limbs resulted in greater magnitude induced changes in MT (red) in the lower limb area of the left posterior limb of the internal capsule (light blue) and the left cerebral peduncle (dark blue).

**Fig. 6:**
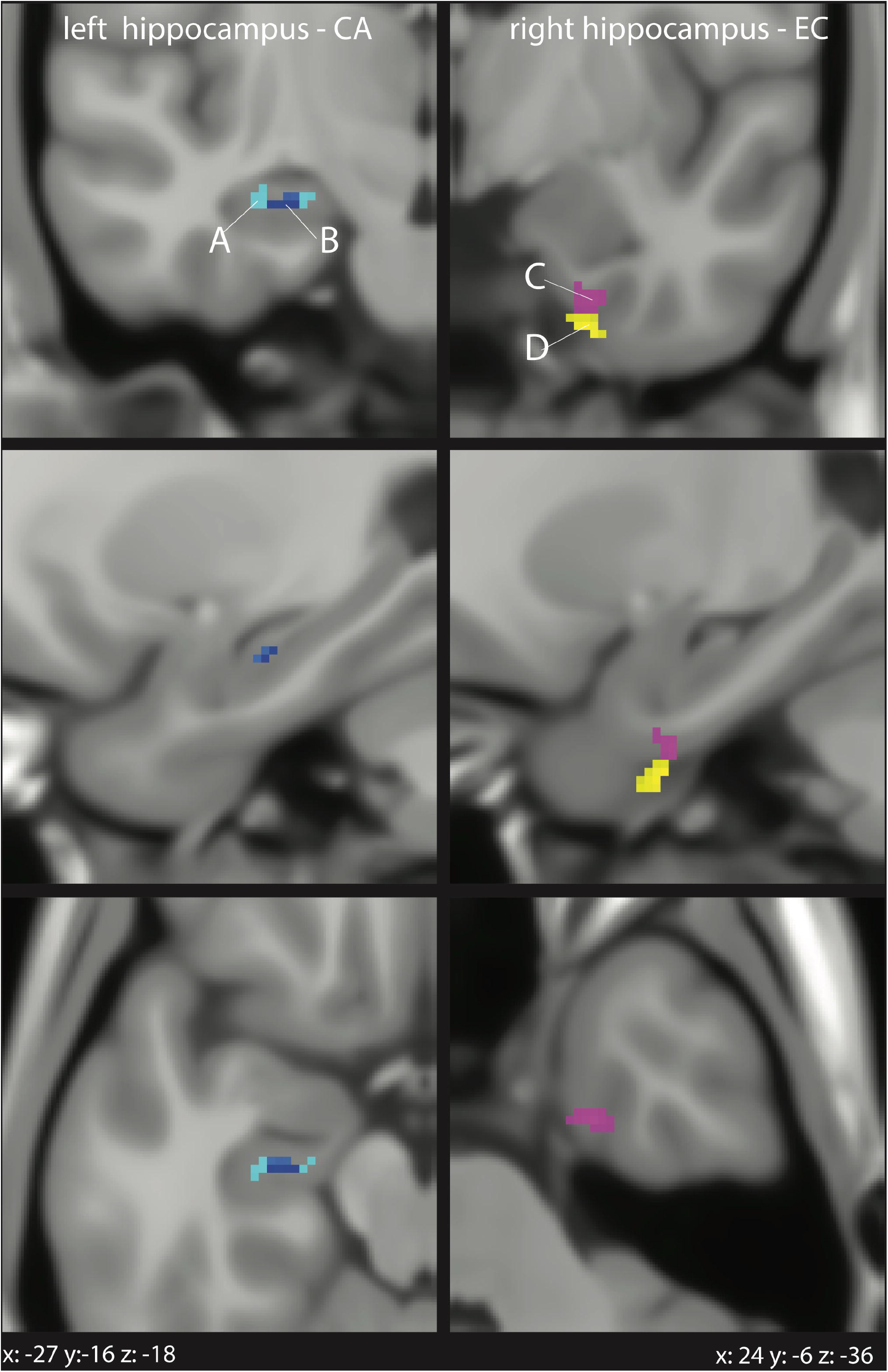
Associations between macro- and microstructural changes and training improvements. In the left hippocampus-CA, negative correlation between quadratic macrostructural changes and improvement in the percentage of correct stimuli responses (A) and a positive correlation between quadratic macrostructural changes and response time improvement (B) were observed. In the right EC, positive correlations between linear MT increases and faster response time improvement (*γ*) (C) as well as between linear MT increases and faster improvement (*γ*) in the percentage of correct stimulus responses (D) were seen.

#### The hippocampus-EC formation

The hippocampus is capable of plastic change in response to tasks—reflecting its diverse roles—with size increases associated with motor learning in mice (Badea et al., 2019) and acquired expertise in human navigation, dancing and juggling in seminal volumetric GM analyses (Boyke et al., 2008; Hüfner et al., 2011). We observed multimodal, dynamic responses to training in the hippocampal-EC formation. A key element of performance improvement is the translation of complex cue patterns into optimal motor response sequences, which are based on experience of the task and a degree of mnemonic sequence learning. The left anterior hippocampus is active in early and later implicit and explicit motor sequence learning (Gheysen et al., 2010; Long et al., 2018) and in offline reprocessing of prior learning (Albouy et al., 2008; Pinsard et al., 2019), which is enhanced by sleep (Pinsard et al., 2019). In patients with right medial temporal lobe epilepsy, better performance in a motor sequence tapping task was associated with larger anterior hippocampal volume. The transient decreases in R1 and MT that we observed in the anterior and dorsal hippocampus, respectively, were accompanied by a similarly transient volume increase in the left anterior hippocampus, which may also be explained by activity-induced angiogenesis and oligodendrogenesis in these GM regions or by classical Hebbian propagation, stabilisation and elimination of dendritic spines (Moore et al., 2017). Conversely, an increase in R1 and MT was observed in the bilateral EC, which provides dense inputs to the hippocampus and is necessary for the acquisition of associative memories during visuomotor learning in monkeys (Yang et al., 2014). These progressive, linear increases in R1 and MT maps may represent myelin modulation subtending task performance through increases in conduction velocities in this region; correlative cellular experiments would be informative.

In the behavioural analysis, performance improvement in Stepmania was accompanied by transient volumetric increase in the hippocampal CA a key site for the consolidation of motor sequence learning (Albouy et al., 2008; Yasuda et al., 2011), while the rapidity of this skill acquisition was subtended by MT contrast increases in the right EC known to have an important role in temporal, spatial and long-term memory (Brun et al., 2008; Suh et al., 2011). These findings may be a manifestation of the dorsal hippocampus’ role as a novelty detector, comparing stored memories in the dentate gyrus and anterior hippocampus with a stream of direct sensory representations from the EC (Duncan et al., 2012). Baseline %CSR was negatively correlated with quadratic MT changes in the EC, indicating that poorer task performance may allow more ceiling for both behavioural improvement and its microstructural proxy in the EC.

Aerobic exercise has been shown to influence hippocampal morphology in animals (Van Praag et al., 2005) and human neuroimaging studies (Firth et al., 2018; Frodl et al., 2019). Exercise generally appears to exert its effect on hippocampal volume in a linear fashion over 6-12 months (Firth et al., 2018), with this effect most pronounced in elderly individuals where exercise drives retention of volume in the context of ongoing, age-related neuronal loss (Firth et al., 2018). The rapid, parabolic changes observed in this study correlated with task mastery and are unlikely to be driven by aerobic exercise alone, although a contribution cannot be excluded.

### Temporal dynamics of multimodal changes across the corticospinal system and the hippocampus

Acquiring five serial scans per subject afforded the opportunity to assess temporal associations between macro- and microstructural changes in brain areas constituting the cortico-limbic networks involved in task learning. Three key interdependencies were revealed in the R1 microstructural data. Firstly, transient R1 changes in the cerebral peduncles preceded those in the left M1, suggesting training-induced microstructural plasticity in the CST can precipitate similar processes in the cortex, either directly or as part of a rostral propagation of microstructural changes over time. The process that these changes represent is unclear, but similar dynamics in these very different GM and WM structures once more implies processes common to both, such as vascular changes or astrocytic remodelling; although, intracortical myelin is abundant in the precentral gyrus (Rowley et al., 2015), so direct myelin effects may be plausible. Secondly, transient decreases in R1 in the M1 and hippocampal CA are linked to later, linear increases of R1 in the EC. This is in keeping with the role of the EC as the interface between the hippocampus and cortex, with the latter sharing limited direct connections (Burman, 2019; Maller et al., 2019). Contemporaneous volumetric increases have been observed in the sensorimotor cortex and the hippocampus during 5 days of visuo-motor training (Kodama et al., 2018), but this is the first evidence of an M1-EC-hippocampal circuit active during motor learning and underlines the potential versatility of quantitative MRI markers in mapping skill acquisition and recovery over time.

### Limitations

While increasingly accepted as a proxy for CNS microstructure and well-correlated with histological myelin markers (Clayden et al., 2015; Natu et al., 2019), R1, R2* and MT remain indirect estimates of microstructure. However, we were able to consistently demonstrate coherent changes consistent with the current understanding of the processes underpinning complex motor learning tasks (Lungu et al., 2014). Selecting an optimal training task for such a study is a compromise, and we faced the additional difficulty of requiring a task that was relevant and challenging, yet achievable by patients with neurological deficits and healthy participants alike. The training duration of four weeks was chosen based on prior studies (Draganski et al., 2006; Makino et al., 2016) and this choice was validated by the fact that almost all participants achieved asymptotic performance in the third or final week of training. The data gathered here will be used in future studies in comparison to patients with neurological injury learning Stepmania. It is for this reason that participant allocation was not fully randomised and instead age-matched to ensure comparability with patients with spinal cord injury. While this may allow confounders such as scanner properties, body weight etc. to influence the results, we believe these are effectively minimised through the use of an exclusively male, righthanded population and intra-scanner coefficient of variation for MPM parameters on our scanner has been determined to be <5% (Leutritz et al., 2020).

Finally, analyses in this study were limited to linear and quadratic temporal changes, although clearly other temporal patterns are possible. The quadratic model has been previously utilised (Wenger et al., 2016) and is particularly suitable for the detection of expansion and renormalisation processes in the brain during learning (Draganski et al., 2006; Hopkins et al., 2018; Moraud et al., 2016). More complex models, including latent change score models (Kievit et al., 2018) or dynamical systems will be explored in the future (Ziegler et al., 2017).

## Conclusion

Serial quantitative MRI parameters were acquired in healthy individuals learning to master the motion game Stepmania using either their upper or lower limbs. Analysis revealed multimodal changes across the somatosensory, corticospinal and hippocampal systems in trainees. In the CST, quadratic R1 and R2* trajectories followed a convex course during training, implying an underlying cellular process characterised by expansion and renormalisation. Somatotopic differences in the magnitude of changes in myelin-sensitive MT were observed between the upper and lower limb training groups in the left CST, suggesting a somatotopy of learning, whereby activity dependent myelination produces faster conduction velocities. The left hippocampus emerged as a focus of multimodal responses to motor learning, with transient volumetric, R1 and MT changes paralleled by linear R1 and MT changes in the bilateral EC, with transient increases in left hippocampal volume and linear changes in EC microstructure associated with skill level and speed of acquisition, respectively. Even more strikingly, R1 microstructural changes in one region consistently preceded those in another, with evidence of a coherent and choreographed network, encompassing the CST, M1 and hippocampal-EC formation active during motor skill learning in healthy individuals. Taken together, these results offer the first insights into the co-ordinated plasticity of a cortico-limbic network, accessible with non-invasive MRI, subtending skill acquisition in the undamaged brain.

## Acknowledgements

We would like to thank all participants for participation in this study. We thank Eric Reese (https://github.com/kyzentun) for selflessly offering his time and expertise in the writing of the Stepmania scripts. We also thank Prof Bogdan Draganski, Dr. Chris Easthope Awai, Dr. Marc Bolliger and Prof. Armin Curt for their guidance and support in developing and carrying out this study; and thanks to Daniel R. Altmann for the statistical support.

